# Inhibition of nonsense-mediated decay in TDP-43 deficient neurons reveals novel cryptic exons

**DOI:** 10.1101/2025.06.28.661837

**Authors:** Irika R. Sinha, Yingzhi Ye, Yini Li, Parker S. Sandal, Philip C. Wong, Shuying Sun, Jonathan P. Ling

## Abstract

TAR DNA-binding protein 43 kDa (TDP-43) is an essential splicing repressor whose loss of function underlies the pathophysiology of amyotrophic lateral sclerosis and frontotemporal dementia (ALS-FTD). Nuclear clearance of TDP-43 disrupts its function and leads to the inclusion of aberrant cryptic exons. These cryptic exons frequently introduce premature termination codons resulting in the degradation of affected transcripts through nonsense-mediated mRNA decay (NMD). Conventional RNA sequencing approaches thus may fail to detect cryptic exons that are efficiently degraded by NMD, precluding identification of potential therapeutic targets. We generated a comprehensive set of neuronal targets of TDP-43 in human iPSC-derived i^3^Neurons (i^3^N) by combining TDP-43 knockdown with inhibition of multiple factors essential for NMD, revealing novel cryptic targets. We then restored expression of selected NMD targets in TDP-43 deficient i^3^Ns and determined which genes improved neuronal viability. Our findings highlight the role of NMD in masking cryptic splicing events and identify novel potential therapeutic targets for TDP-43-related neurodegenerative disorders.

## Introduction

TAR DNA-binding protein (*TARDBP*, TDP-43) is a ubiquitously expressed, predominantly nuclear protein implicated in a variety of diseases^1–3^ and a major component of ubiquitinated cytoplasmic inclusions in amyotrophic lateral sclerosis and frontotemporal dementia (ALS-FTD)^4–6^. Missense mutations in *TARDBP*, the gene encoding TDP-43, established the role of TDP-43 in the pathogenesis of ALS-FTD^7–10^ and the possibility of a toxic gain-of-function mechanism underlying ∼97% and ∼45% of post-mortem ALS and FTD cases, respectively^11,12^. Conversely, studies from human disease suggest TDP-43 loss-of-function as the early driver of neuron loss in ALS-FTD^13,14^ as well as other diseases with TDP-43 pathology, such as limbic-predominant age-related TDP-43 encephalopathy (LATE)^15–17^, inclusion body myositis (IBM)^18–20^, and Alzheimer’s disease^17,21–27^.

TDP-43 plays a key role in transcriptional and translational fidelity by regulating cryptic splicing. It preferentially binds to uracil-guanine repeats ([UG]_n_) in pre-mRNA^28–31^ and prevents the inclusion of nonconserved cryptic exons (CEs) into mature transcripts^14,32–34^. When TDP-43 function is lost, the aberrant inclusion of CEs frequently introduces premature termination codons (PTCs) and other mRNA disruptions including alternative polyadenylation (polyA), alternative transcriptional start sites, altered untranslated regions (UTR), or in-frame insertions^35,36^. Importantly, in-frame and early polyA CE are more likely to be expressed at measurable levels in patient tissue and biofluids^16,17,24,26,27,37–45^. Conversely, excessive TDP-43 expression can lead to the skipping of constitutive exons^10,46^. The inclusion of CEs is often cell-type specific and not conserved across species^14,47–49^. These mRNA changes disrupt RNA and protein availability, with PTCs often triggering nonsense-mediated mRNA decay (NMD) that degrades the aberrant mRNA prior to translation and can lead to reduced protein expression levels ^50–53^. To fully understand and treat the cellular effects of TDP-43 dysfunction, it is crucial to identify its targets and the processes to which they contribute.

RNA-sequencing (RNA-Seq) provides comprehensive transcriptome coverage and can be used to detect alternative splicing isoforms, including CEs. Alignment software for RNA-Seq reads can detect canonical and noncanonical splice junctions via split reads^54^. The output of the aligner can be parsed by splicing analysis tools and be used to identify potential novel junctions of significance within a condition^55–57^. Finally, visualizers, such as UCSC Genome Browser^58^, and wet lab experiments can be used to confirm the CEs. These techniques, when previously applied to experimental models and patient post-mortem samples, have been used to identify CEs associated with TDP-43 depletion^36^. However, most previously identified CEs are detected due to inefficient NMD processing (Figure 1A). As NMD efficiency is both transcript and cell type dependent^59^, CEs that undergo efficient NMD likely remain undetected in many datasets.

**Figure 1.**
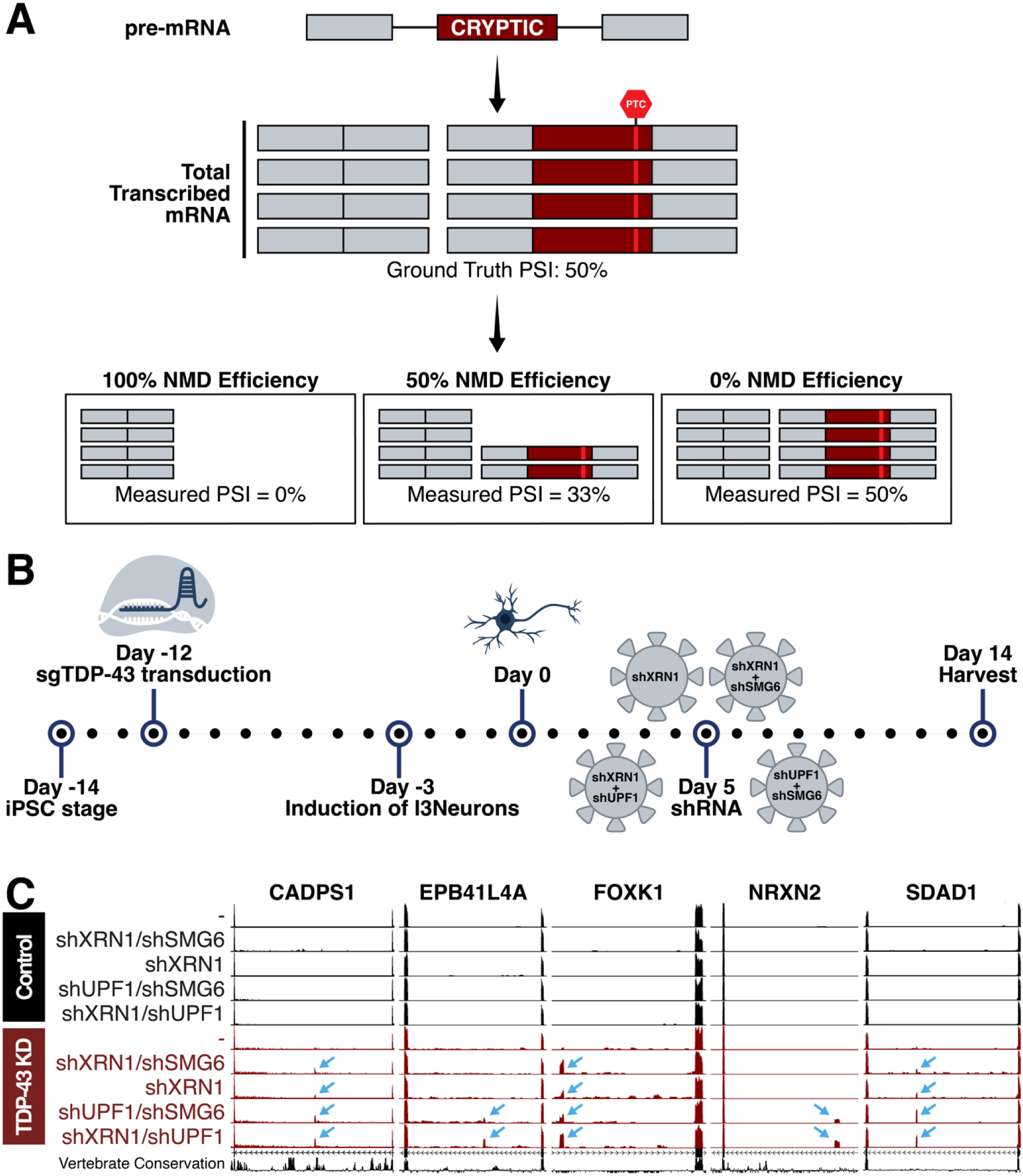
Nonsense-mediated decay (NMD) was inhibited to identify TDP-43 gene targets which undergo efficient degradation. (**A**) Pre-mRNA transcribed in a cell includes cryptic exons when TDP-43 is dysfunctional. This pre-mRNA is processed into mRNA and the cryptic exons often encode PTCs that are targeted by NMD. Efficiency of NMD impacts measured cryptic exon PSI after RNA sequencing or RT-PCR. (**B**) Schematic of experimental timeline. i^3^Ns with a CRISPRi module were transduced with sgTDP-43 12 days prior to differentiation to create a stable TDP-43 knockdown cell line. Five days post-differentiation, we conducted shRNA-mediated knockdown of NMD factors *XRN1*, *UPF1*, and *SMG6*. i^3^Ns were harvested 14 days after differentiation and RNA extracted for RNA sequencing. (**C**) Cryptic exons in *CADPS1, EPB41L4A, FOXK1, NRXN2,* and *SDAD1* are measurable by RNA sequencing after NMDi. These cryptic exons are not identifiable in cells with only TDP-43 knockdown or NMDi and are specific to TDP-43 knockdown conditions.

Cells utilize NMD as a quality control mechanism to prevent the translation of abnormal or undesired protein products^50–52,60,61^. NMD factors are recruited and act on transcripts containing PTCs and lead to the elimination of unproductive transcripts^62–66^. Although NMD does not act equally on all targets, CE-containing transcripts that are efficiently destroyed by NMD remain undetectable with methods such as RT-PCR and RNA-Seq. Consequently, these cryptic targets are largely absent from previously compiled CE studies.

To identify TDP-43 targets and to create a dictionary of CEs in iPSC-derived i^3^Neurons^67–69^ (i^3^Ns), we depleted TDP-43 from i^3^Ns in combination with NMD factors *XRN1*, *SMG6*, and/or *UPF1* and performed RNA-Seq on the i^3^Ns. Inhibition of each of these factors alone is known to hamper NMD processes and we found that combined depletion of *UPF1* and *XRN1* led to the greatest detection of otherwise hidden CEs. Altogether, we identified 421 unique TDP-43-associated i^3^N CEs in 329 genes. From these CEs, we selected genes known to be associated with other disorders to selectively knock down in i3Ns and found no single knockdown to recapitulate the loss of cell viability in i3Ns after TDP-43 knockdown. Finally, we conducted rescue experiments in TDP-43 depleted i^3^Ns and found overexpression of some targets, such as *ZDHHC11*, moderately increased cellular viability but no single target fully rescued toxicity.

This new dictionary of i^3^N TDP-43 cryptic targets with information such as location, canonical junctions, and type provides researchers an accessible reference without needing to re-align and re-analyze existing sequencing data. Furthermore, these targets can be used as readouts of TDP-43 depletion in i^3^Ns and other cell-types that incorporate them after TDP-43 depletion. Additionally, our knockdown and rescue experiments demonstrate the importance of rescuing TDP-43 targets as a whole rather than rescuing single genes. These findings have significant implications for future therapeutic strategies designed to restore TDP-43 splicing capabilities.

## Results

### NMD inhibition (NMDi) in TDP-43 deficient i^3^Ns uncovers a novel set of cryptic targets

To model TDP-43 dysfunction in cortical neurons, we knocked down TDP-43 in i^3^Ns^67–69^, a well-studied human induced pluripotent stem cell-derived cell line^41,44,70–76^ (Figure 1B, Supplementary Figure 1A). As CEs can introduce premature termination codons (PTCs) in mRNA and therefore likely trigger nonsense-mediated decay (NMD) of these aberrant transcripts, we depleted NMD factors in conjunction to TDP-43 depletion using shRNA-mediated knockdown to inhibit degradation of CE-associated transcripts (Figure 1B, Supplementary Figure 1A-B). By preventing the degradation of CE-including targets, we increased the percent-spliced-in (PSI) values of the CEs, improving our ability to measure them^77^. To inhibit NMD, we used shRNA-mediated knockdown of key NMD factors. Previous work have shown that the combinatorial knockdown of some factors, such as *XRN1* with *UPF1*^63,77^ or *SMG6*^51^ improve efficacy of NMDi. We thus tested the following conditions: sh*XRN1*, sh*XRN1 +* sh*UPF1*, sh*XRN1* + sh*SMG6*, and sh*UPF1* + sh*SMG6* (Figure 1A). *XRN1* knockdown conditions had a general upregulation of *SMG6* and the combinatorial knockdowns of sh*XRN1 +* sh*UPF1* and sh*UPF1* + sh*SMG6* were most successful at knocking down target genes (Figure 1C, Supplementary Figure 1B, E-F).

The i^3^Ns were harvested 14 days post-differentiation and sequenced using bulk RNA-Seq. Differential splicing analysis was used to identify CEs included because TDP-43 depletion, and PSI values were calculated for the cryptic junctions. Identified differential splicing events were confirmed as TDP-43 targets if a [UG]_n_ motif was present near the CE. Instances of complex splicing, such as the inclusion of multiple CEs in the same canonical junction (Supplementary Figure 1C), were evaluated on a case-by-case basis.

From the i3N data, we identified 421 TDP-43-associated CEs, some of which have been previously described^14,34,37,38,40,41,44,49^ (Table 1). These CEs were categorized as cassette, exon extension, early polyA, or alternate start site CEs (Figure 2A, Supplementary Figure 2A). 5.5% of CEs were considered leaky with an average cryptic junction PSI (avgPSI) of greater than 20% in at least one control condition, 8.9% generated in-frame peptides in the final protein product, and 7% were exons already annotated by GENCODE but present only at low levels or not present in tissues sequenced for the Genotype-Tissue Expression (GTEx) project^78,79^. Additionally, chromosome 19 was enriched in cryptic exons (Supplementary Figure 1D), although its density of AC/TG repeats is similar to the other chromosomes^80^. Skiptic splicing events^10,46^, the skipping of canonical exon, can occur as a consequence of excessive TDP-43 overexpression and were therefore excluded from these analyses (e.g. *POLDIP3*^81^, *KCNQ2*^26,44,49^). We additionally investigated which targets were most impacted as a result of TDP-43 depletion and termed those incorporated with an avgPSI of at least 70% in any TDP-43 knockdown condition and an avgPSI less than 10% in any control condition (Supplementary Table 1, Supplementary Figure 2B, 2D). These CEs, we termed as “high expressor” CEs^80^.

**Figure 2.**
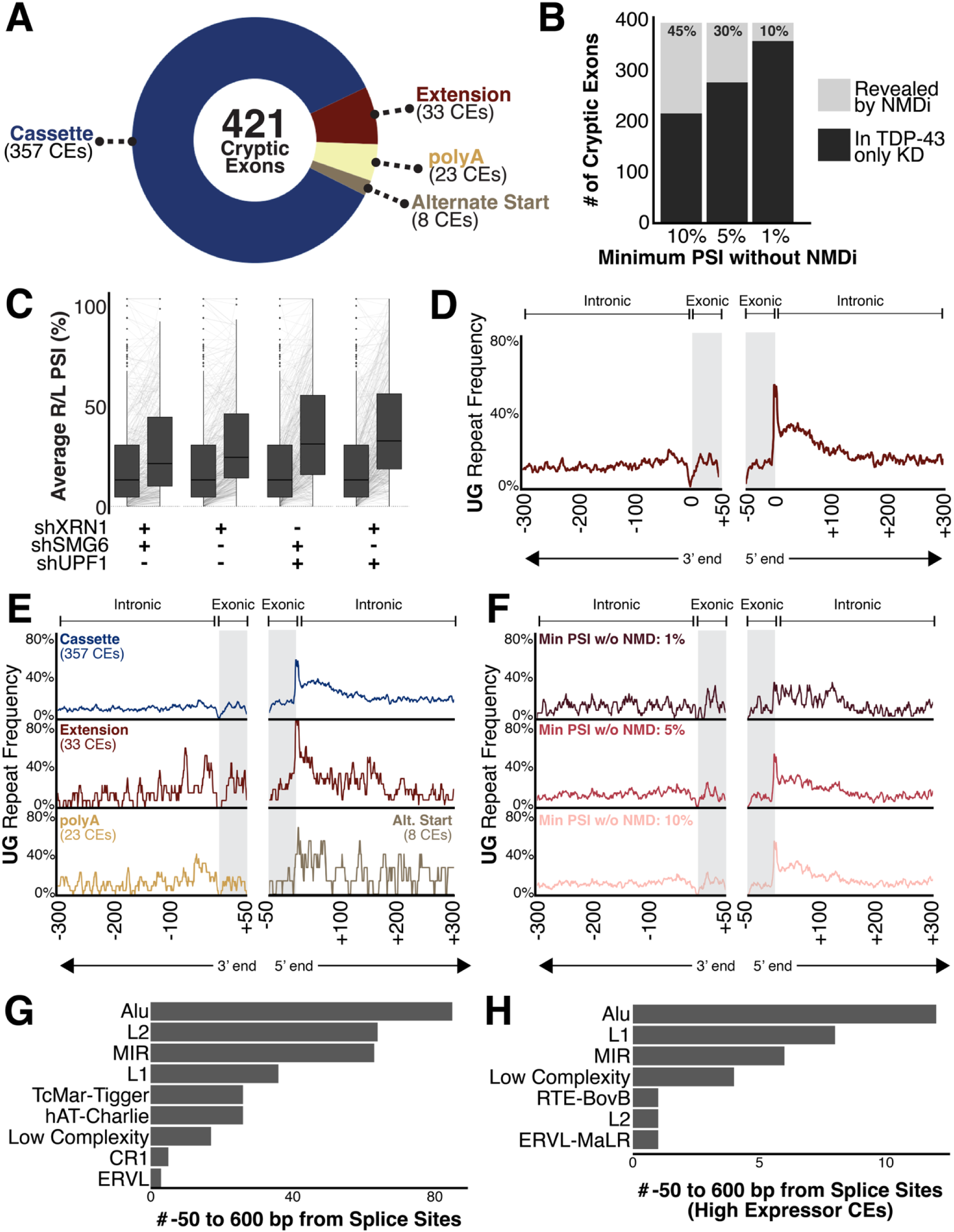
(**A**) 421 total CEs were identified in i^3^Ns and categorized as cassette exons, exon extensions, early polyadenylation events, or alternative start sites. 26 CEs were “leaky” in the control conditions. (**B**) Comparison of CEs measurable with and without NMDi based on given cut-offs. Each cut-off defines the minimum avgPSI needed for a CE to be considered measurable without NMDi. (**C**) Comparison of median cryptic exon avgPSIs between conditions. (**D-F**) Frequency of [UG]_n_ dinucleotide repeats near CE splice sites. TDP-43 CEs often have [UG]_n_ after the 3’ end of the CE. (**D**) Full compendium of 421 CEs. (**E**) Split by type of CE (cassette, exon extension, polyA, alternative splice site). Exon extension CEs are enriched for [UG]_n_ repeats after the 3’ end of the cryptic exon compared to the overall average and other types. PolyA cryptic exons have a slight enrichment for the [UG]_n_ before the 5’ end of the cryptic exon. Due to lack of 3’ and 5’ exon-exon junctions for polyA and alternative start site CEs, respectively, they only have the repeat mapped near one splice site. (**F**) Split by measurability without NMDi. Cut-offs are the same as in Figure 2C. CEs with PSI less than 1% in TDP-43 knockdown conditions have less pronounced [UG]_n_ repeats at the 3’ end of the CE. (**G**) UCSC RepeatMasker annotations present at least twice between 600 bp external each splice site and 50 bp internal of the sites. Alu family repeats are most common near cryptic exons. (**H**) UCSC RepeatMasker annotations present at least twice between 600 bp external each splice site and 50 bp internal of the sites of high expressor CEs. Alu family repeats are most still common.

Finally, to maximize the comprehensive nature of this compendium, we cross-referenced the cryptic exons in our list with those identified in other studies using a TDP-43 knockdown model in i^3^Ns^44,49^. Our model included most CEs found in other studies although the CEs in two genes, *HS6ST3* and *MGAT5B*, were originally screened out due to a lack of reads which include the canonical exon-exon splice junction. Additionally, it appears that while the CE in *HS6ST3* is included at high levels after TDP-43 knockdown, the gene itself is not normally expressed well in i^3^Ns. We also include in this compendium well-known CEs such as the one in *AGRN*, even though they are not measurable in this cell-type.

### UPF1 knockdown produced the most robust NMDi in TDP-43 deficient i^3^Ns

Of these 421 CEs, many were either ambiguous or even unmeasurable prior to NMDi (Figure 1C). Some genes with well-known CEs, such as *EPB41L4A*, were discovered to have additional CEs, often incorporated at a different rate, while several other new gene targets were identified. To determine which CEs required NMDi for identification, we set three different possible thresholds for CE inclusion in the TDP-43-KD-only i^3^Ns: 1%, 5%, and 10% (Figure 2B). At a threshold of 1%, CEs measurable in at least 1% of sequenced transcripts for a gene are identifiable without NMDi. At this most strict threshold, 10% of CEs are considered “hidden” because of NMD and only measurable in conditions with NMDi (Figure 2B, Supplementary Figure 2C). This indicates that 90% of these CEs are inefficiently degraded by NMD and could be measured in other instances of TDP-43 dysregulation. Additionally, these findings confirm that the downregulation of many genes in TDP-43-associated disorders is in part attributable to CE-driven degradation.

Of the different NMDi conditions, the one which led to the greatest median increase in CE detection was the combinatorial inhibition of *XRN1* and *UPF1*, although the median increase due to the combinatorial inhibition of *UPF1* and *SMG6* was not significantly different (Figure 2C, Supplementary Figure 1E-F). In part, this is likely due to *UPF1*’s role as crucial component of NMD initiation and target recognition^52,61,64,66^. One of several pathways through which *UPF1* initiates NMD involves the recruitment of *SMG6* to cleave target mRNAs by phosphorylated *UPF1* and the subsequent degradation of the 3’ cleavage product by the exonuclease *XRN1*^52^. Additionally, recent findings indicate TDP-43 depletion in motor neurons disrupts *UPF1* phosphorylation and, consequently, *UPF1*-mediated pathways such as NMD target recognition^82^. As a result, it is likely the direct knockdown of *UPF1* via shRNA was compounded by the indirect knockdown due to TDP-43 depletion. This NMDi was then further strengthened by co-depletion of other NMD factors.

### TDP-43 binding [UG]_n_ motif enriched near the 3’ end of cryptic exons

For each identified CE, we mapped the location of UG repeats proximal to either side of each splice site. The splice site of the 3’ end of the CE is the donor site for the following, canonical exon. This donor site is generally UG in the pre-mRNA and there is a relative spike in UG motif in that area. Almost 60% of all identified CEs include that donor site in the nearby UG repeat that is likely the binding site for TDP-43.

For each identified CE, we mapped and compared the location of UG dinucleotide repeats between 50 bp within the CE to 600 bp external of each cryptic splice site. Consistent with previous reports^83^, we found a relative enrichment of [UG]_n_ repeats near this site and, notably, nearly 60% of the identified CEs include this site as part of the proximal [UG]_n_ repeat (Figure 2D). This is consistent across both cassette and exon extension CEs, along with other types of CEs, such as noncoding and in-frame (Figure 2E, Supplementary Figure 2E).

High-expressing CEs have a slightly greater enrichment and length, on average, of the [UG]_n_ repeats at the 3’ end of the CE (Supplementary Figure 2F). Both exon extension and early polyA CEs also have an increased enrichment of repeats near the 5’ end of the CE as compared to the overall compendium (Figure 2E). CEs uncovered by NMDi are also enriched for a 3’ [UG]_n_ repeat at the 5% and 10% cut-offs but the pattern is far less distinct at the 1% cut-off, indicating a potentially weaker binding site for those least observed CEs (Figure 2F). Additionally, all the CE events have no enrichment of other tested dinucleotide repeats such as [CA]_n_, [CG]_n_, [AA]_n_, or [UU]_n_ (Supplementary Figure 2G), which are associated with other RNA-binding proteins such as FUS^84^. Beyond simple repeats, such as the dinucleotide repeat, we also searched the UCSC RepeatMasker database for other families of repeats proximal to the CEs. Of these, Alu family repeats were most enriched near the CEs near high-expressing CEs (Figure 2G-H).

### TDP-43-associated cryptic exon-containing genes are highly impacted by NMD

Combining TDP-43 knockdown with NMDi is crucial for understanding the true impact of TDP-43 depletion on transcriptome. In addition to unmasking CEs hidden by NMD, we are also able to better identify genes most affected by the loss of TDP-43. By quantifying the actual percentage of mature transcripts which incorporate each CE, we better understand the full scope of pathways and cellular processes impacted by TDP-43 dysfunction. The median avgPSI for CEs increases from 12.7% after TDP-43 knockdown alone to 31.3% with co-depletion of *XRN1* and *UPF1* (Figure 2C, Supplementary Figure 1E-F). This indicates a large percentage of CE-including transcripts are being depleted by NMD prior to translation. High expressor CE targets also display increase in avgPSI after NMDi, although the overall incorporation of these CEs are relatively high in all conditions (Supplementary Figure 2D).

To identify genes highly impacted by TDP-43 depletion, we subset our CEs to the top 30 high expressor CE targets, as ranked by maximum avgPSI in TDP-43 knockdown conditions, include well-known CEs such as those in *CAMK2B, FBN3,* and *PFKP* (Figure 3A, Supplementary Figure 2B). Some CEs, such as the one in *ADCY8*, are sequenced at low levels after TDP-43 knockdown. Initial results suggest that only 30% of *ADCY8* mRNA incorporate the CE, although NMDi reveals that the CE is incorporated in close to 85% of transcripts. Therefore, the top gene targets identified by TDP-43 knockdown alone are not sufficient for understanding the most affected genes and that NMDi is necessary for determining the most important targets. This highlights that the top gene targets identified through TDP-43 knockdown alone do not represent the most affected genes, as NMD inhibition reveals a broader set of cryptic exon inclusions that would otherwise be missed.

**Figure 3.**
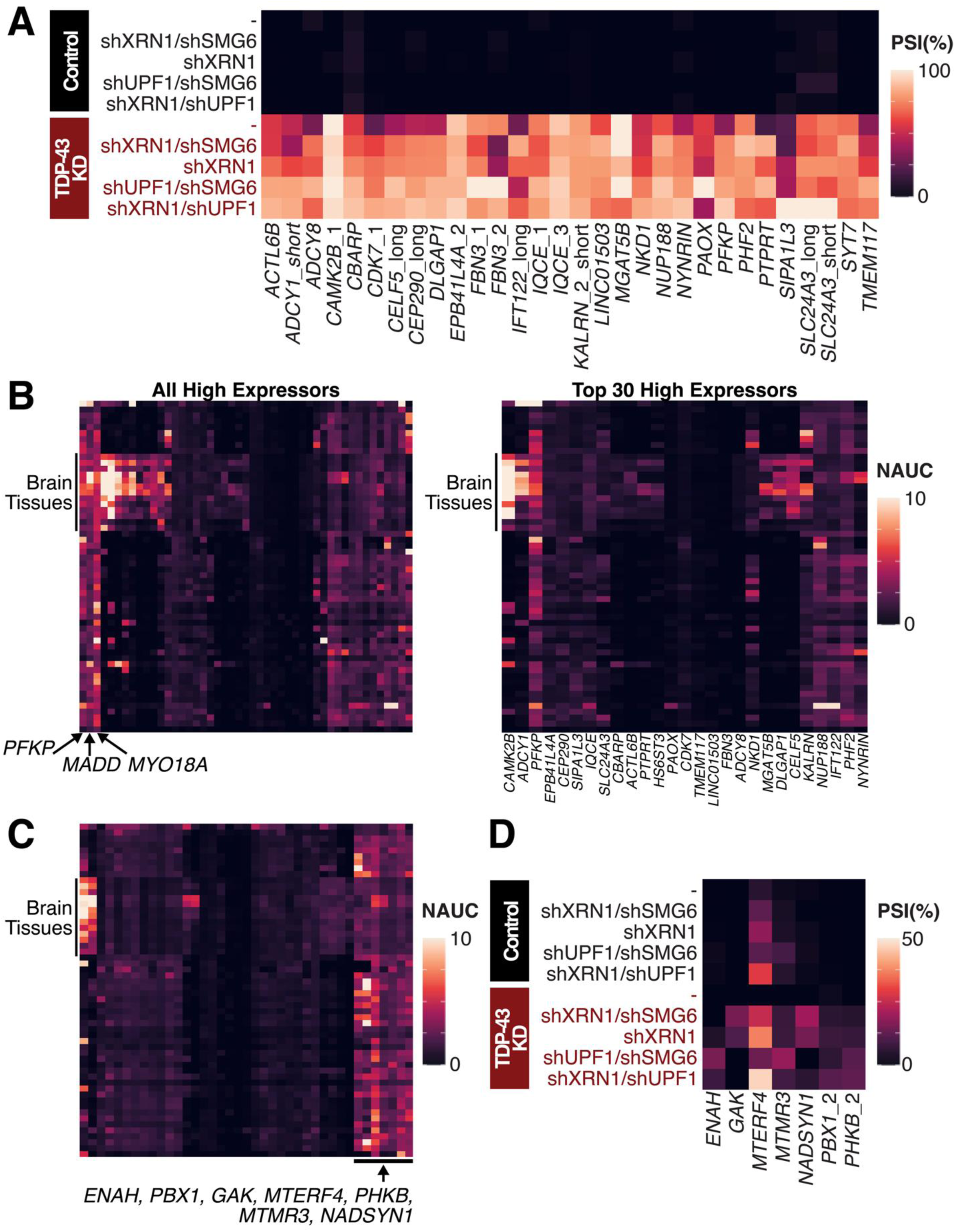
PSI and Gene Expression Levels of Cryptic Exons. (**A**) avgPSI in different conditions of top 30 cryptic exon gene targets, as ranked by maximum avgPSI in any TDP-43 knockdown condition. Many targets are transcribed at a higher frequency in at least one NMDi condition than expected from non-NMDi TDP-43 knockdown condition. (**B-C**) Gene expression levels as measured by normalized area under the curve (NAUC) extracted from ASCOT^79^ for high expressor CEs. Different tissues are represented by rows and different CEs by the columns. (**B**) (*Left*) All high expressor CEs (*Right*) Top 30 CE gene targets as per 3A. (**C**) All CEs with less than 1% PSI after only TDP-43 knockdown. (**D**) CEs with less than 1% PSI after only TDP-43 knockdown and ubiquitous expression across different tissue types.

CE-containing genes can be potential biomarkers for ALS-FTD. One criterion for biomarkers is for measurable expression in tissues of interest. For the identified CEs, we investigated the tissue-specific enrichment patterns as measured in GTEx samples through ASCOT (Figure 3B, Supplementary Figure 3). Some high expressor CE gene targets, such as *PFKP, MADD,* and *MYO18A* have ubiquitous expression across tissues and are likely to have measurable levels of NMD-associated depletion after TDP-43 dysfunction – potentially in non-brain associated tissue. Others, such as *CAMK2B,* are enriched in brain tissues and are more likely to be measured in biofluids such as CSF or post-mortem as a readout for TDP-43 loss of function. Further investigation of the CE genes discovered by NMD also pinpoints *ENAH, PBX1, GAK, MTERF4, PHKB,* and *MTMR3* as ubiquitously expressed with measurable levels of cryptic exons (Figure 3C).

### Overlap between known ALS-associated genes and TDP-43 targets

While only 10% of ALS-FTD cases occur in a familial or genetic pattern, numerous genes have been linked to ALS-FTD with associated mutations or through correlation studies^85,86^. Of the 40+ genes identified as ALS-FTD-associated, our study confirms three as direct targets of TDP-43 splicing regulation: *ATXN1*, *CHMP2B,* and *UNC13A*.

The recent identification of the cryptic exon in *UNC13A*^40,41^, and its adjacent ALS-linked risk variant, has led to the development of isoform-correcting antisense oligonucleotide therapeutic trials^36,76^. By combining NMD inhibition with TDP-43 knockdown, we were also able to find CEs in both *ATXN1* and *CHMP2B* (Figure 4A). Both CEs are incorporated at low levels, if at all, after TDP-43 knockdown only. The CE in *CHMP2B* is measurable with TDP-43 knockdown alone but much clearer after NMDi. It is also incorporated at low levels in TDP-43 depleted SH-SY5Y cells^41^. However, in our TDP-43 depleted i^3^Ns, we were unable to detect the CE in *ATXN1* without NMDi; it is additionally not included in *ATXN1* after TDP-43 knockdown in HeLa cells^14^, SH-SY5Y cells^41^, or iPSC-derived motor neurons^37^. This indicates that at least part of the genomic dysregulation of these genes after TDP-43 knockdown is due to direct effects of cryptic exon inclusion and transcript degradation rather than downregulation at the transcriptional level.

**Figure 4.**
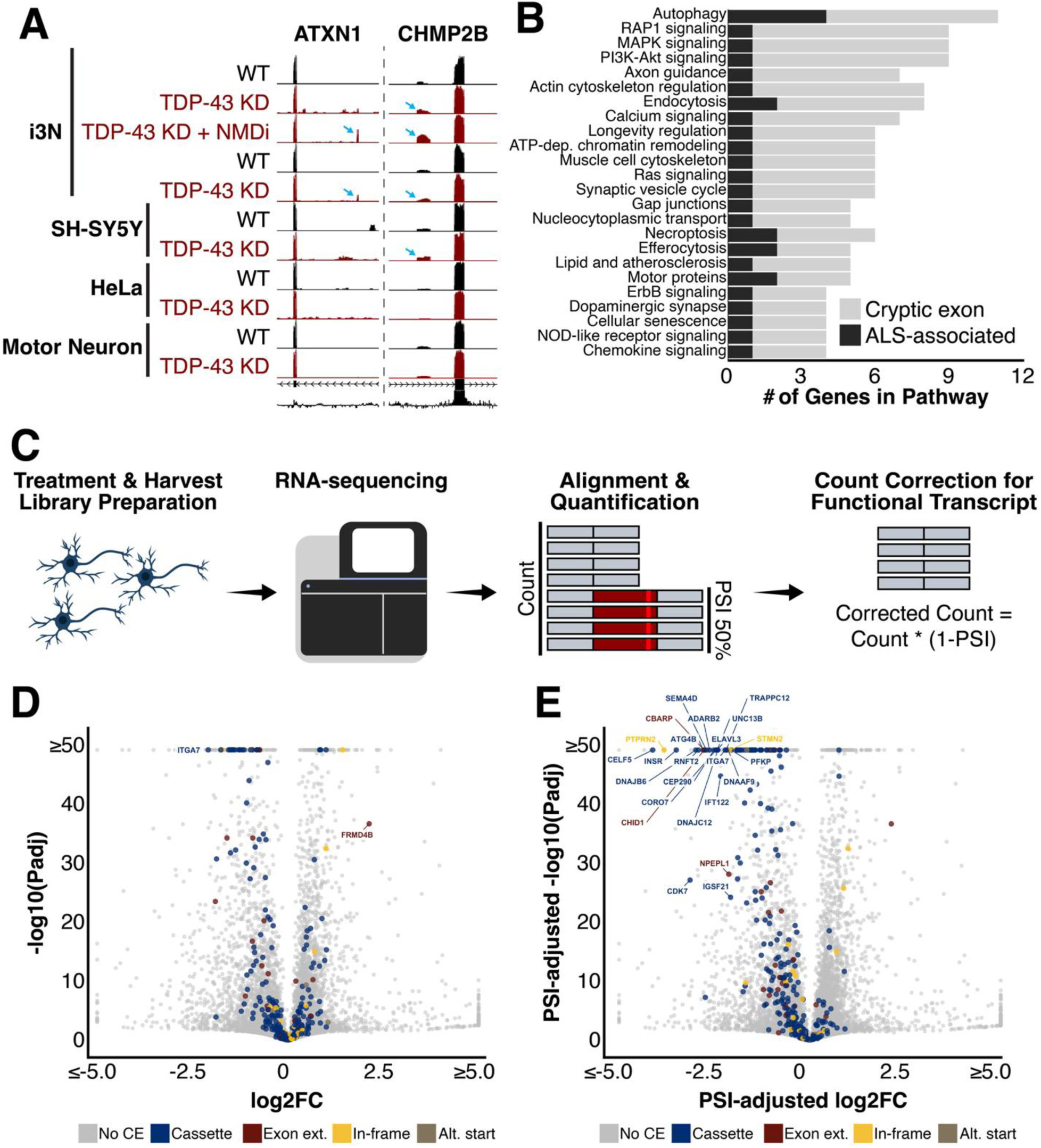
Connections to ALS-FTD and differential gene expression. (**A**) *ATXN1* and *CHMP2B* are ALS-associated risk genes that contain cryptic exons (**B**) Biological pathways including multiple ALS-associated risk genes and cryptic exon-containing genes include autophagy (**C-E**) Differential gene expression after TDP-43 knockdown of cryptic exon-containing genes (**C**) Schematic describing transcript count correction for functional transcripts. I3N were treated and harvested. Prepared RNA library was sequenced and STAR and Salmon were used to align and quantify transcripts. Transcript counts for CE-including transcripts were corrected for functional gene expression by multiplying total count by the percentage of transcripts without CE. (**D**) Normal method without any adjustment (**E)** Transcript counts adjusted for average PSI of cryptic exon which would decrease the number of transcripts encoding functional protein

In addition to genes directly impacted by TDP-43-associated splicing dysregulation, we also investigated cellular pathways which included both ALS-FTD-associated genes and TDP-43-associated CEs. Patients carrying a mutation in an ALS-FTD-associated gene with TDP-43 depletion may be more susceptible to disruptions in these cellular processes. Of the tested pathways which included at least one ALS-FTD-associated gene and at least one CE gene, the most impacted pathway appeared to be the autophagy pathway (Figure 4B). Several additional pathways contained both ALS-FTD-associated genes and genes with TDP-43-associated CEs, suggesting functional overlap and potential avenues of further exploring the consequences of TDP-43 dysfunction.

### TDP-43-associated cryptic exons increase functional downregulation of target genes

Alongside the splicing analysis of the i^3^Ns, we conducted differential gene expression analysis of the sequenced RNA (Figure 4C). As expected, the expression of several genes is dysregulated as a result of TDP-43 knockdown. Of the CE-including genes, 24 have significant expression changes of at least two-fold. These include downregulated genes such as *STMN2*, *CORO7*, and *CAMK2B*. Only two CE-including genes, *ITGA7* and *FRMD4B*, have significant changes in gene expression greater than four-fold (Figure 4D). *ITGA7* is downregulated while *FRMD4B* is upregulated, potentially to compensate for a decrease in functional protein availability since 61% of transcripts incorporate an out-of-frame CE after the knockdown of TDP-43. For these CE-including genes, it is also important to note that at least part of the downregulation is a result of increased CE-related transcript degradation rather than transcriptional regulation. It is even possible that at least some of these are transcriptionally upregulated to compensate for increased degradation rates.

Another consideration regarding gene expression levels is that transcript quantification tools do not distinguish between functional and non-functional transcripts. As a result, the true extent of functional gene loss cannot be understood without adjusting for reduced levels of functional transcripts. To better understand the impact of TDP-43-associated splicing dysregulation on gene expression, we adjusted the total transcript counts for CE-including genes to account for this inclusion (Figure 4C). After doing so, the number of significantly downregulated genes with expression fold-changes greater than 4 increased dramatically (Figure 4E).

### No single gene target knockdown recapitulates TDP-43 knockdown toxicity

Of the 421 CE-including genes, we selected 56 genes with known disease relevance and essential gene functions for a cellular toxicity screen. This subset included genes with in-frame CEs due to the possibility of the CE interrupting a functional protein domain. Each gene was depleted in i^3^Ns via shRNA-mediated gene knockdown, without TDP-43 depletion. Most shRNAs were validated by the vendor but, for some genes, two different shRNAs were tested due to the lack of prior validation. The resultant toxicity was then quantified using propidium iodide (PI) staining (Figure 5A). PI is a fluorescent DNA stain impermeant to live cells and used to mark dead cells. Control shRNA treatment resulted in minimal toxicity, with little PI signal colocalizing with DAPI-stained nuclei. In contrast, shRNA knockdown of TDP-43 led to extensive PI and DAPI colocalization, showing widespread cell death and reflecting the highly toxic nature of the knockdown.

**Figure 5.**
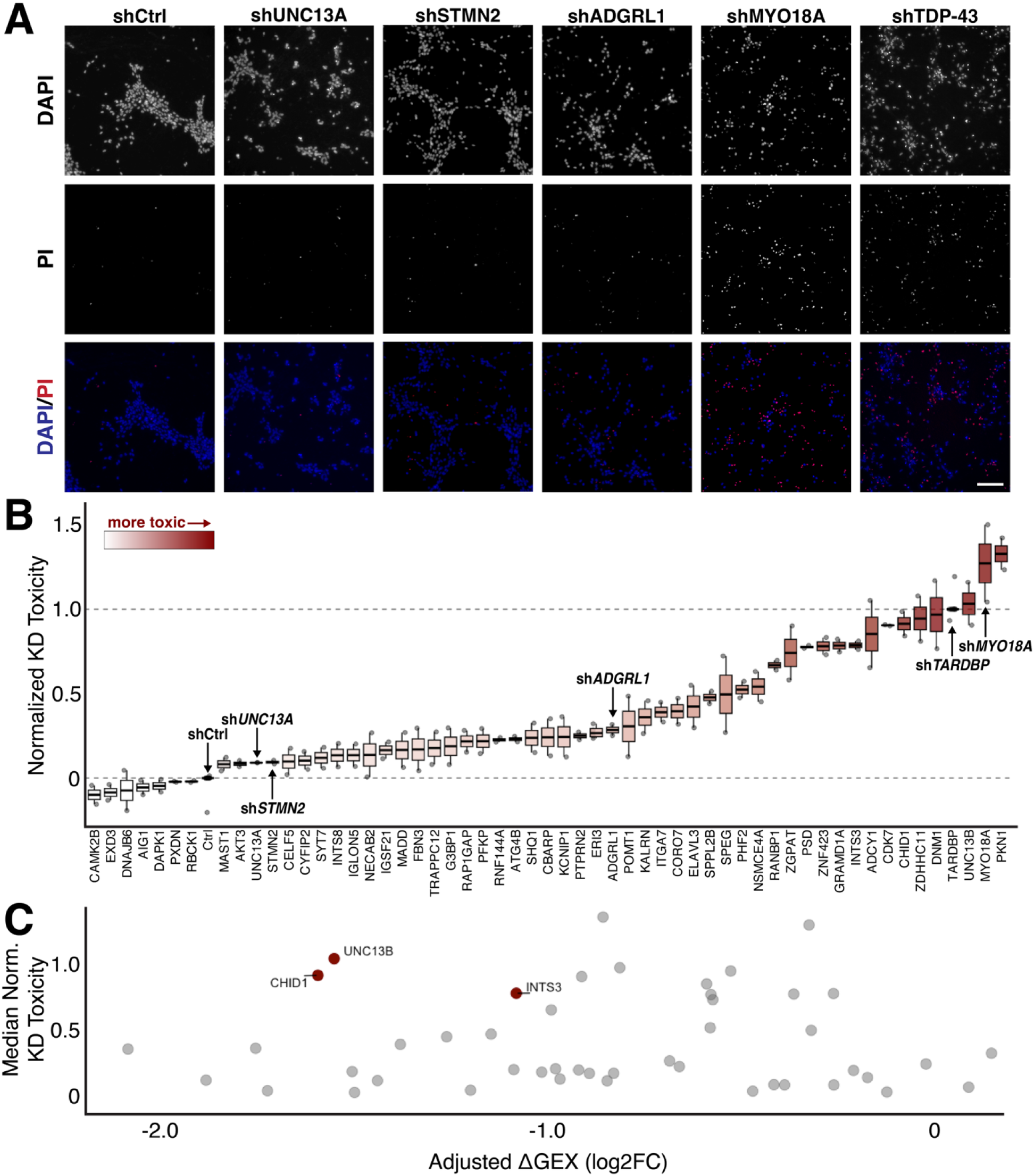
shRNA-mediated knockdown of cryptic exon-containing genes. (**A**) Sample images of shRNA-treated i^3^Ns co-stained for DAPI and propidium iodide (PI), which stains dying cells; shCtrl is a non-targeting control. Scale bar = 200µm. (**B**) Ranked quantification of shRNA toxicity normalized to shCtrl and shTDP-43. Min-max normalization was used with maximum toxicity set to shTDP-43 and minimum toxicity to shCtrl. Most shRNA treatment leads to intermediate toxicity and does not recapitulate TDP-43 toxicity. (**C**) Comparison of cryptic exon gene adjusted downregulation in i^3^Ns after TDP-43 knockdown with toxicity of shRNA-mediated knockdown. Genes with at least a two-fold downregulation and toxicity at least 50% of shTDP-43 are highlighted and labeled.

We then used CellProfiler to identify and quantify nuclei and PI staining within each image. Then, we determined the percentage of dead cells associated with each gene knockdown and normalized the resultant toxicity to that measured for TDP-43 knockdown and transduction of a control shRNA (Figure 5B). Therefore, a toxicity score of 0 indicates similar PI+ cell percentage to the control while a toxicity score of 1 indicates a similar PI+ cell percentage to TDP-43 knockdown. For targets with multiple tested shRNA, the construct with greater toxicity was used in the analysis.

Knockdown of several genes alone, such as *MYO18A* and *ZDHHC11*, led to toxicity comparable to TDP-43 knockdown alone (Figure 5B). However, most knockdowns led to a cell death phenotype in-between the control and TDP-43 knockdown. Several conditions had no toxicity at all. This indicates that cell death as a result of TDP-43 knockdown is likely due to a combination of mechanistic failures rather than the dysregulation of a single target after splicing-associated degradation and downregulation.

The significance of the difference in median knockdown toxicity from the control condition was then assessed. We compared knockdown toxicity with the effective downregulation as a result of CE inclusion (Figure 4E, Figure 5C). The only genes with high knockdown toxicity that were also highly downregulated were *CHID1*, *UNC13B*, and *INTS3*. Since the CE in *UNC13B* is both in-frame and somewhat leaky^73^ - hence identifiable in people without TDP-43 deficits - it is likely that this CE actually does not decrease functional transcripts. This makes *CHID1* and *INTS3* better candidates for rescue in a TDP-43 knockdown.

### TDP-43 knockdown toxicity cannot be rescued by a single target

Should TDP-43-related toxicity and neuronal death be primarily driven by CE inclusion in one or a few specific genes, then restoring those targets might be sufficient to rescue the toxic effects. Based on CellProfiler-derived toxicity scores and manual screening of the most toxic knockdowns, we selected 15 candidate genes for rescue experiments in TDP-43–depleted i3Ns (Figure 6A). As noted, two CE-containing genes are downregulated by over 50% following TDP-43 depletion and induce cellular toxicity when individually knocked down, and these were included in the final list (Figure 5C). shRNA-mediated knockdown of TDP-43 was conducted on day 4 following differentiation of i^3^Ns and lentiviral overexpression was conducted on day 7. The cells were then imaged on day 14 and cell survival was quantified manually using PI staining.

**Figure 6.**
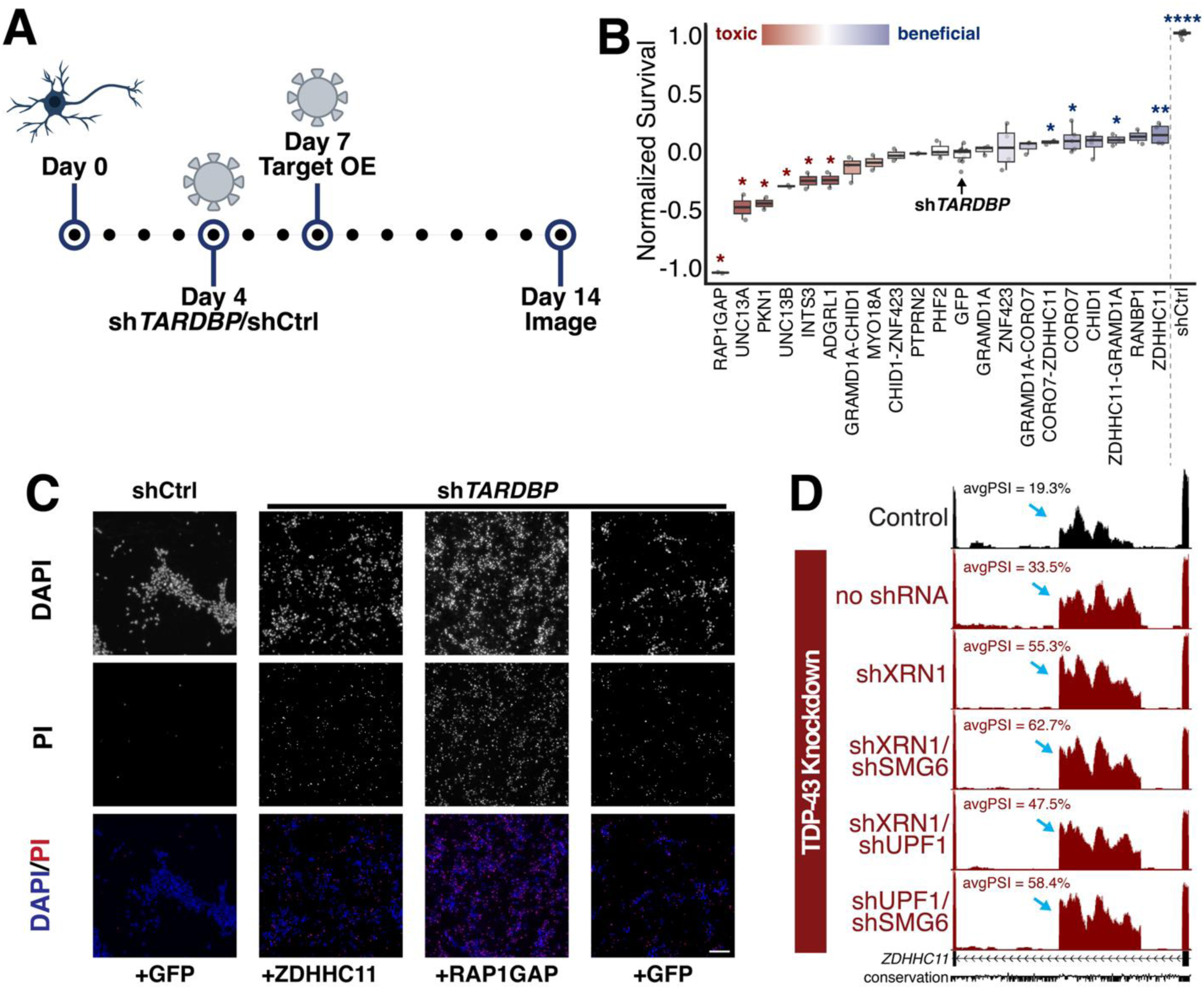
Rescue experiments in i^3^Ns with TDP-43 knockdown. (**A**) Schematic of rescue experiments in i^3^N where cells are transduced with shTDP-43 or shCtrl (non-targeting control) on day 4 post-differentiation and then target genes are overexpressed on day 7 post-differentiation. (**B**) Quantification and ranking of normalized cell toxicity after over-expression of genes after shTDP-43 treatment. Min-max normalization was used with minimum survival set to shTDP-43 and maximum to shCtrl. Values less than 0 correspond with decreased survival compared to shTDP-43 with GFP overexpression and greater than zero correspond with increased survival. Conditions with survival significantly different from shTDP-43+GFP treatment are marked. (**C**) Sample images from rescue experiments of negative control (shCtrl with GFP overexpression) and three conditions of shTDP-43-treated i^3^N. *ZDHHC11* overexpression led to the highest survival in comparison to shTDP-43 with GFP overexpression while *RAP1GAP* overexpression led to the lowest. Scale bar = 100µm. (**D**) *ZDHHC11* cryptic exon is leaky in the control but increased significantly after TDP-43 knockdown.

Similar to the analysis for the single gene knockdown experiments, cell survival was normalized to the shControl + GFP overexpression and the shTDP-43 + GFP overexpression conditions. A survival score of 0 indicates similar survival to cells with TDP-43 depletion while a survival score of 1 indicates similar survival to the control cells.

Of all the overexpressed genes, no condition was sufficient to bring cell survival to the same level as the control cells (Figure 6B). Some overexpressed genes, such as *RAP1GAP*, caused increased toxicity in the cells after overexpression (Figure 6B-C). *INTS3* and *CHID1* overexpression did not lead to any significant increase in survival when compared to TDP-43-depleted cells either. Only cells treated with *CORO7 + ZDHHC11, CORO7, GRAMD1A + ZDHHC11,* and *ZDHHC11* overexpression displayed a significant increase in survival. Although some conditions combined the overexpression of *ZDHHC11* with other genes, it was the single overexpression of this gene that appeared to lead to the best survival in TDP-43 depleted cells (Figure 4B-C). This is interesting as, although the gene is associated with important functions as a palmitoyltransferase and in immune responses, it is also one of the genes with a leaky cryptic exon. 19.3% of transcripts in control i^3^Ns also incorporate this cryptic exon, potentially as a mechanism of RNA regulation. However, after TDP-43 knockdown, this increases to 33.5% and NMDi raises it even further to 58.4% (Figure 6D). This drastic increase in cryptic exon incorporation and the nearby [UG]_n_ repeat confirmed its status as a TDP-43-dependent cryptic exon. Altogether, our data finds single gene rescues through overexpression are insufficient to rescue TDP-43-associated toxicity and suggests a combinatorial approach or CE-directed therapeutics for increased efficacy.

## Discussion

Over the past decade, researchers have systematically catalogued TDP-43 cryptic exons across various cell types and experimental conditions. However, conventional RNA-Seq analyses have largely captured only those CEs that escape robust NMD, thereby underrepresenting the full scope of TDP-43 dysregulation. In this study, we leverage co-depletion of TDP-43 and NMD decay factors to generate a comprehensive list of CEs normally targeted directly in i^3^Ns by TDP-43 splicing regulation in i^3^Ns, including those that are otherwise undetectable under normal conditions. NMD efficiency varies across cell types and contexts^59^, thus extending these inhibition studies to other cell types beyond neurons will be essential for understanding the systemic impact of TDP-43 dysfunction.

This curated CE dataset can be used as a reference for mining public RNA-Seq data to detect TDP-43 loss-of-function and to better understand the pathways disrupted in disorders such as ALS-FTD and Alzheimer’s disease. For each CE, we include extensive annotations including genomic coordinates, mean PSI values for each experimental condition, reading frame status, and associated gene. We hope this resource will help facilitate future work that uncovers disease-associated SNPs that may have been overlooked in datasets generated without NMDi. These data may also drive future biomarker discovery efforts and guide therapeutic target prioritization.

Our study indicates that TDP-43 loss-of-function influences more genes and pathways than previously understood, including direct impacts on two ALS-associated genes, *ATXN1* and *CHMP2B*. The relationship between these genes and TDP-43 in ALS-FTD is complex, however, given that ATXN1 mutations typically confer gain-of-function effects, whereas the CHMP2B cryptic exon is already partially incorporated when NMD is inhibited even without TDP-43 loss. Conversely, when NMDi does not efficiently degrade cryptic exons after TDP-43 knockdown, the true impact on gene expression is underestimated given that many reads originate from aberrant, non-functional isoforms. By integrating PSI values into gene expression calculations, we can more precisely quantify the levels of functional transcripts.

While generating this list of CEs, we also ran a parallel screen to investigate the toxicity of single TDP-43 target knockdowns with the toxicity of TDP-43 depletion itself. We then overexpressed the most promising targets in TDP-43 depleted i^3^Ns to test whether any gene could independently rescue the toxicity of TDP-43 knockdown. Results from both the knockdown screen and overexpression assays indicated that TDP-43 toxicity does not arise from one target, but from the cumulative effect of all genes and pathways impacted by TDP-43 depletion.

A key limitation of our screens is that shRNA knockdown of cryptic targets may drive gene expression lower than it falls after TDP-43 loss, whereas overexpression of target genes may push gene expression above physiological levels. Nevertheless, our results show that overexpression of certain genes containing TDP-43 CEs, such as *UNC13A*, may worsen toxicity. This suggests that therapeutic efforts should remain focused on antisense oligonucleotide (ASO) as opposed to gene therapies that deliver the gene. Partial restoration of other CE-containing genes, while not achieving cell viability, may also achieve functional benefits.

Our work underscores the need for therapeutic approaches that restore TDP-43’s function as a splicing regulator to effectively address the underlying pathology. When TDP-43 is depleted, neurons accumulate cryptic exons that destabilize key transcripts and ultimately lead to cell death. While restoring a single target may be beneficial^36,87^, correcting the full spectrum of TDP-43 dysregulation will likely require rescuing multiple genes in concert. Approaches that address this underlying dysfunction hold great promise for treating TDP-43-associated disorders.

## Commonly used acronyms/terms

**ALS**: Amyotrophic Lateral Sclerosis

**avgPSI**: Average value of the PSI of both the right and the left junction of a cryptic exon (if cassette). For non-cassette cryptic exons, this simply refers to the cryptic junction PSI.

**CE**: Cryptic exon

**FTD**: Frontotemporal Dementia

**High expressor CE**: CE with a PSI of 70% or greater in at least one TDP-43 knockdown condition and 10% or less in all control conditions

**I^3^Ns**: I^3^Neurons – human induced pluripotent stem cell-derived cell line with cortical phenotype

**NMD**: nonsense-mediated decay

**NMDi**: NMD inhibition

**PI**: Propidium iodide – a stain impermeant to living cells used to quantify cell death

**polyA**: Alternative polyadenylation

**PSI**: Percent spliced-in value of an exon-exon junction in mRNA

**TDP-43**: Protein encoded by *TARDBP* gene with splicing repressor functions. Associated with ALS-FTD.

**UTR**: Untranslated region

## Methods

### Human Induced Pluripotent Stem Cell (iPSC)-derived i^3^Neuron (i^3^N) Cell Culture

Human i^3^N iPSCs were maintained in Essential 8 media (ThermoFisher #A1517001) on plates coated with Matrigel (Corning #354277) and differentiated into i^3^N on Day 14 post-induction as previously detailed^67–69^. All cells were maintained at 37°C with 5% CO_2_.

### Stable Knockdown of TDP-43 in i^3^N using Clustered Regularly Interspaced Short Palindromic Repeat Inactivation (CRISPRi)

To achieve a stable knockdown of TDP-43, i^3^N-iPSCs^67^ with a constitutively expressed CRISPRi module^69^ were transduced with a lentivirus expressing an sgRNA targeting TDP-43 between the 1st and 2nd exons and prior to the translational start site (seq: GGCCCGCGCGTGCCAGCCGA). The stable iPSC line was then established as previously described^39^.

### NMD Inhibition in sgTDP-43 i^3^Neurons

sgTDP43 i^3^Neurons at differentiation day 5 were transduced with lentivirus expressing shRNAs targeting XRN1, UPF1, and/or SMG6 (Supplementary Table 2) at 3 MOI individually or at 3 MOI each in combination. The neurons were harvested on differentiation day 14.

### Comparison of Cryptic Target and ALS-associated Gene Pathways

The clusterProfiler package^88,89^ was used to identify pathways from the Kyoto Encyclopedia of Genes and Genomes (KEGG) containing both cryptic target genes and ALS-associated genes ^85,86^.

### RNA Sequencing

RNA was extracted from cells in trizol and Stranded mRNA Prep Kit (Illumina) was used to prepare libraries as per the manufacturer’s protocol. Sequencing was performed using an Illumina NextSeq 500 to generate 150bp paired-end reads.

### RNA Sequencing Analysis

Nextflow nf-core RNAseq (v3.18.0-gb96a753)^90,91^ pipeline built with Nextflow (v24.10.5, build 5935) was used to process FASTQ files using default presets. Briefly, FastQC (v0.12.1) was used to check read quality, Trim Galore! (v0.6.10) was used for adapter trimming, STAR (v2.7.11b) was used for alignment to GRCh38, and Salmon (v1.10.3) was used to quantify transcripts. Finally, bedtools (v2.31.1) was used to generate bigWigs and DESeq2 (v1.28.0)^92,93^ was used to perform differential gene expression analysis. Data were visualized on the UCSC Genome Browser (http://genome.ucsc.edu/)^58,94,95^.

Pipeline output DESeqDataSet object was used to visualize differential gene expression differences using R between control and sgTDP43 i^3^N. Raw counts data were exported from DESeqDataSet and adjusted for functional gene expression by the following formula:

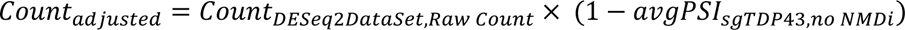

Using adjusted counts dataframe, a new DESeq2 object was created and re-calculated differential gene expression. R scripts were used to visualize gene expression fold change and compare to toxicity of shRNA knockdowns.

Splicing analysis of the datasets were conducted in multiple ways to increase thoroughness. First, annotation-free identification of alternative splicing events were identified using a “junction-walking” method as previously described^79^. Next, we used FRASER (v2.0)^57,96^ and Leafcutter (v0.2.9)^56^ with default settings to find any potentially missed or low level alternative splicing patterns in the data. Cryptic splice junctions identified were visualized on the UCSC Genome Browser and screened using the following criteria: 1) Splice junction is not present in normal controls or has a PSI at least 20% greater than control, 2) [UG]_n_ repeat present within 600 bp of either cryptic exon splice site, 3) At least one splice site is present in STAR output .sj files. Skyptic exon events were excluded from this analysis but some potential events, along with previously described events, are available in Supplementary Table 2.

Computational resources from the Advanced Research Computing at Hopkins (ARCH) were used to run nf-core RNAseq, FRASER, and LeafCutter pipelines.

### shRNA Generation

shRNA constructs for the non-targeting control (shCtrl) and TDP-43 (shTDP-43) were generated following the Addgene protocol for modifying the RNAi Consortium pLKO.1 cloning vector (addgene.org/protocols/plko/). shCtrl contains a non-hairpin insert as recommended by the RNAi consortium (Addgene #10879). In brief, shTDP-43 was generated by cutting the pLKO.1 cloning vector (Addgene #10878) with AgeI (NEB # R0552) and EcoRI (NEB #R0101) and then ligated with the target sequence as per Supplementary Table 3. All other shRNAs (Supplementary Table 3) were synthesized by MilliporeSigma.

### Gene Knockdown Toxicity Assays

i^3^N on differentiation day 4 were transduced with lentivirus expressing shRNA against target genes at 1 MOI individually. An RNAi Consortium-recommended non-targeting control shRNA (shCtrl) was used as a negative control and generated as previously described^70,97^. On day 14, neurons were stained with Hoechst 33342 (Cell Signaling Technology #4082S) and Propidium Iodide (ThermoFisher #P1304MP) for 15 minutes and then imaged. Live cells were determined by Hoechst staining and dead cells were determined by co-stain of Hoechest and propidium iodide staining. CellProfiler (v4.2.8) was used to locate nuclei and propidium iodide specks and then a custom R script was used to calculate percentage of propidium iodide positive nuclei (pctPI+) and the normalized toxicity of each shRNA. Normalized toxicity was calculated using Min-max normalization to scale values to a 0 to 1 range. Given that Toxicity_min_ is the median pctPI+ of the shCtrl of the same plate and Toxicity_max_ is the median pctPI+ of the shTDP43 of the same plate:

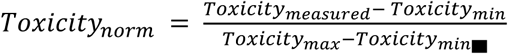

CellProfiler pipeline details can be found on GitHub. The toxicity of each knockdown was ranked and then screened manually. The most promising 15 target genes were selected for the overexpression rescue experiment.

### TDP-43 Toxicity Rescue Assays

TDP-43-targeting shRNA or shCtrl was added to i^3^N on differentiation day 4 and lentiviral transduction was used on differentiation day 7 to overexpress each selected target gene. On day 14, i^3^N were stained with Hoechst 33342 (Cell Signaling Technology #4082S) and Propidium Iodide (ThermoFisher #P1304MP) for 15 minutes and then imaged. Manual counting was used to determine live and dead cell counts using Hoechst stain and co-stain of Hoechest and propidium iodide, respectively. Survival was defined as the percentage of propidium iodide negative nuclei (pctPI-). Normalized survival was calculated using Min-max normalization to scale values to a 0 to 1 range. Given that Survival_min_ is the median pctPI-of the shTDP43 + GFP OE of the same plate and Survival_max_ is the median pctPI-of the shCtrl + GFP OE of the same plate:

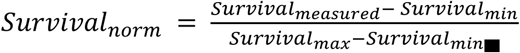

Significant differences in mean survival compared to only a TDP-43 knockdown were determined using the Wilcoxon test.

## Supporting information

Table_Main

## Acknowledgements

This work was supported in part by the Alzheimer’s Association (to J.P.L.), the Institute for Data-Intensive Engineering and Science (to J.P.L.), the NIH RF1NS095969 and RF1NS129878 (to P.C.W.), the RF1NA127925 and RF1AG078948 (to S.S.), ALS Finding a Cure (to P.C.W.), the ALS Association (to P.C.W.), the US Food and Drug Administration (no. 1U01FD008129 to P.C.W.), and Toffler Scholar Award (to Y.Y.).

This material is based upon work supported by the National Science Foundation (NSF) Graduate Research Fellowship Program under Grant No. DGE2139757 (to I.R.S.). Any opinions, findings, and conclusions or recommendations expressed in this material are those of the authors and do not necessarily reflect the views of the NSF.

This work was supported by resources from the Advanced Research Computing at Hopkins (ARCH) core facility (rockfish.jhu.edu), which is supported by the NSF (no. OAC 1920103). We thank Dr. Ricardo de Souza Jacomini for his support installing Leafcutter.

We thank Katherine E. Irwin and Anya A. Kim for their troubleshooting support and suggestions.

## Data Availability

RNA sequencing data for this paper has been deposited in the National Center for Biotechnology Information Sequencing Read Archive (SRA) as BioProject PRJNA1235234. Other RNA-Seq data used in this study are available on the SRA under study numbers ERP126666^41^, SRP057819^14^, and SRP166282^37^.

## Code availability

Scripts for analysis and visualization of data are available in the following GitHub repository: https://github.com/irikas/NMDi_TDP43kd

## Author Contributions

I.R.S., Y.Y., S.S., P.C.W, and J.P.L. conceptualized the study. I.R.S., Y.Y., and J.P.L wrote the manuscript, and all other authors edited and approved. I.R.S., Y.Y., Y.L., and P.S.S. performed experiments and/or analyzed data. I.R.S. and Y.Y. contributed equally and have the right to list their name first in their CV and presentations.

## Notes

### Competing Interest Statement

The authors have declared no competing interest.

