## Supplementary_Figures for "Inhibition of nonsense-mediated decay in TDP-43 deficient neurons reveals novel cryptic exons"

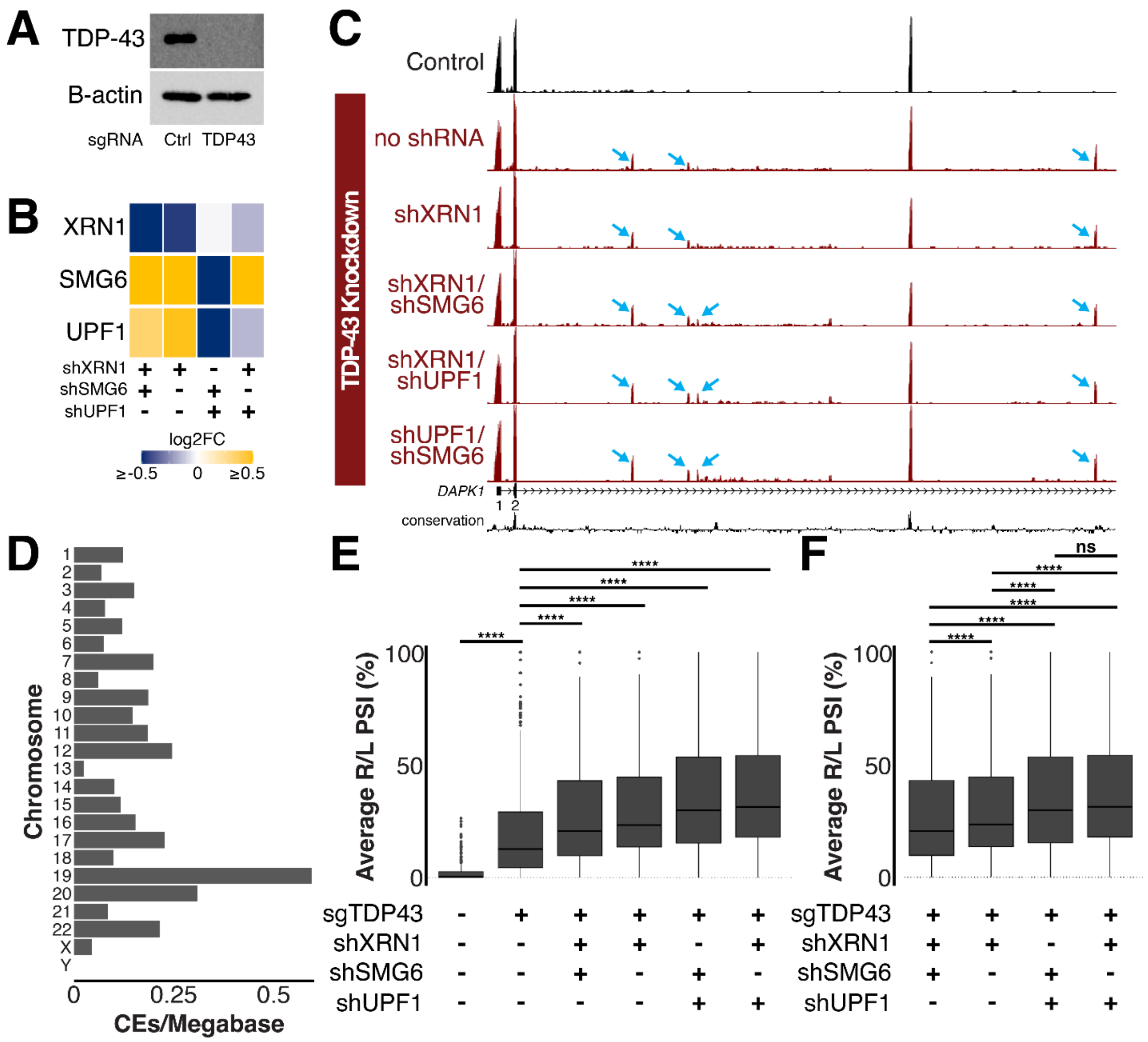


**Supplementary Figure 1. Knockdown of TDP-43 and NMD factors lead to identification of cryptic exons.** (**A**) Immunoblot of TDP-43 after sgTDP-43 treatment shows robust TDP-43 knockdown. (**B**) Differential gene expression of NMD factors *XRN1*, *SMG6*, and *UPF1* after knockdown treatments. (**C**) Four cryptic exons in *DAPK1* gene indicated using blue arrows. All are in the intron between exon 2 and 3. (**D**) Visualization of the number of cryptic exons per megabase of chromosome indicates enrichment of cryptic exons in chromosome 19. (**E**) Average cryptic junction PSI of cryptic exons in control and treated cells. TDP-43 knockdown leads to significant ($p \leq 0.0001$) increase in cryptic junction PSI from control by paired Wilcox test. NMDi conditions further significantly increase ($p \leq 0.0001$) average cryptic junction PSI from TDP-43 knockdown paired Wilcox test. (**F**) Average cryptic junction PSI of cryptic exons in TDP-43 knockdown with NMDi conditions. Mean cryptic junction PSI is not significantly different between shUPF1+shSMG6 and shXRN1+shUPF1 but all other conditions have significantly different mean cryptic junction average PSI ($p \leq 0.0001$) by Wilcox test.

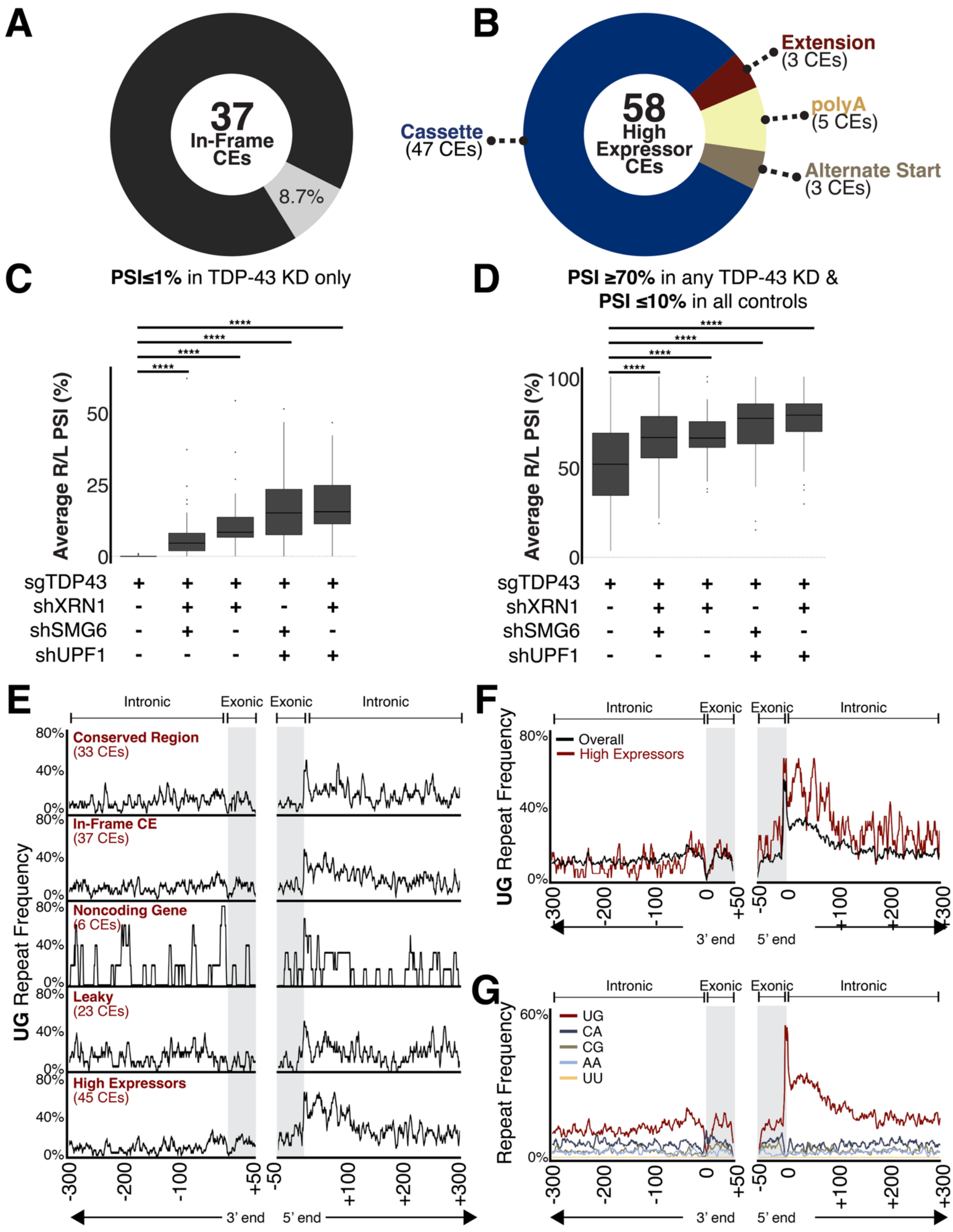

A
37
In-Frame
CEs
8.7%
B
58
High
Expressor
CEs
Extension
(3 CEs)
polyA
(5 CEs)
Alternate Start
(3 CEs)
Cassette
(47 CEs)
C
PSI≤1% in TDP-43 KD only
Average R/L PSI (%)
sgTDP43
shXRN1
shSMG6
shUPF1
D
PSI ≥70% in any TDP-43 KD &
PSI ≤10% in all controls
Average R/L PSI (%)
sgTDP43
shXRN1
shSMG6
shUPF1
E
Intronic
Exonic
Exonic
Intronic
Conserved Region
(33 CEs)
In-Frame CE
(37 CEs)
Noncoding Gene
(6 CEs)
Leaky
(23 CEs)
High Expressors
(45 CEs)
UG Repeat Frequency
3' end
5' end
F
Intronic
Exonic
Exonic
Intronic
Overall
High Expressors
UG Repeat Frequency
3' end
5' end
G
Intronic
Exonic
Exonic
Intronic
UG
CA
CG
AA
UU
Repeat Frequency
3' end
5' end

**Supplementary Figure 2. Knockdown of TDP-43 and NMD factors lead to identification of cryptic exons.** (**A**) 8.7% of identified cryptic exons (CEs) lead to in-frame peptides when translated. (**B**) 58 high expressor CEs were identified with greater than 70% PSI in at least one TDP-43 knockdown condition and less than 10% PSI in all control conditions. Similar to the overall compendium, cassette exons are most common. (**C-D**) Comparison of median cryptic exon avgPSIs between conditions. Significant differences in average PSI of NMDi conditions compared to TDP-43 knockdown only were determined using the Wilcoxon test. (**C**) CEs with less than 1% inclusion in TDP-43 knockdown only condition. (**D**) High expressor CEs. (**E-F**) Frequency of $[UG]_n$ dinucleotide repeats near CE splice sites. (**E**) Split to compare CEs in conserved regions, that encoded in-frame peptides, in noncoding genes, that were leaky in control conditions, and that were high expressors. (**F**) Overlaid comparison of repeat enrichment of overall compendium and high expressors indicate increased enrichment of the repeat at the 3' end of CE. (**G**) Comparison of the frequency of $[UG]_n$, $[CA]_n$, $[CG]_n$, $[AA]_n$, and $[UU]_n$ dinucleotide repeats near CE splice sites.

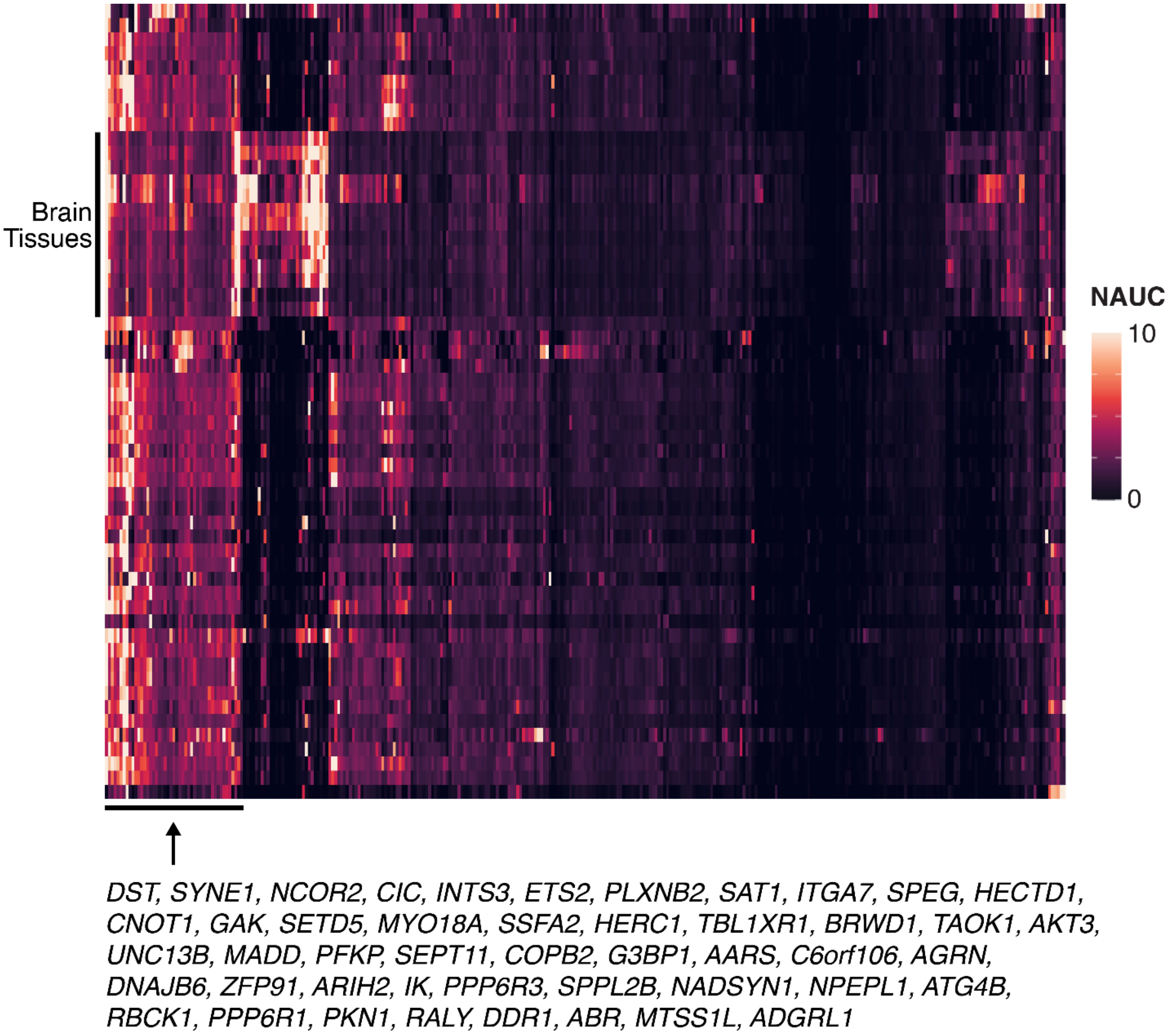


**Supplementary Figure 3. Expression of all CEs across different tissues.** Gene expression levels as measured by normalized area under the curve (NAUC) extracted from ASCOT for all CEs. Different tissues are represented by rows and different CEs by the columns. CE genes with ubiquitous expression across tissues are identified.
